# Phospho-RNA sequencing with CircAID-p-seq

**DOI:** 10.1101/2020.12.16.423093

**Authors:** Alessia Del Piano, Ruggero Barbieri, Michael Schmid, Luciano Brocchieri, Silvia Tornaletti, Claudia Firrito, Luca Minati, Paola Bernabo, Ilaria Signoria, Fabio Lauria, Thomas H. Gillingwater, Gabriella Viero, Massimiliano Clamer

## Abstract

Accurate positional information concerning ribosomes and RNA binding proteins with respect to their transcripts is important to understand the global regulatory network underlying protein and RNA fate in living cells. Most footprinting approaches generate RNA fragments bearing a phosphate or cyclic phosphate groups at their 3′ end. Unfortunately, all current protocols for library preparation rely only on the presence of a 3′ hydroxyl group. Here, we developed circAID-p-seq, a PCR-free library preparation for 3′ phospho-RNA sequencing. We applied circAID-p-seq to ribosome profiling, which produces fragments protected by ribosomes after endonuclease digestion. CircAID-p-seq, combined with the dedicated computational pipeline circAidMe, facilitates accurate, fast, highly efficient and low-cost sequencing of phospho-RNA fragments from eukaryotic cells and tissues. While assessing circAID-p-seq to portray ribosomes engaged with transcripts, we provide a versatile tool to unravel any 3′-phospho RNA molecules.

## Background

RNA molecules bearing a phosphate or cyclic phosphate group at the 3’ end (3′P/2’-3’cP) are generated by either heat fragmentation [1], ribonucleases (e.g. RNase A superfamily) [2], ribozymes [3, 4] or toxins [5, 6]. Beside endogenous 3′P/2’-3’cP RNA molecules [7–11], several biochemical methodologies designed to obtain genome wide positional information of RNA-protein interaction by RNA sequencing require an enzymatic digestion step that generates phosphorylated 3’ termini prior to library preparation. This is the case for near-nucleotide resolution RNA footprinting protocols used to understand the global regulatory network underlying protein and RNA fate in living cells. These protocols are based on (i) RNA:protein cross-linking and immunoprecipitation (CLIP-seq) [12–14] (ii) selective RNA:protein immunoprecipitation (RIP-seq) [15, 16], (iii) RNA:protein affinity purification (uvCLAP) [17], and (iv) ribosome profiling (Ribo-seq) for sequencing of RNA fragments protected by ribosomes. Commonly used enzymes for RNA footprinting include the RNase A superfamily (e.g. RNase I) [18] RNase T1 [19], and RNase T2 [20], which produce 3′ phosphate or cyclic phosphate RNA molecules. Additionally, phosphate or cyclic phosphate group at the 3’ end are also generated by heat fragmentation [21] to define RNA secondary structures. There are currently no technologies to selectively and directly sequence 3′P/2’-3’cP RNA fragments. In fact, the terminal 3′P/2’-3’cP is usually removed by enzymatic reactions before library preparation. This step is likely to result in the loss of useful data. Methods that provide insights into 3′-phospho RNA species rely on an indirect detection of these fragments by means of a periodate treatment [22] and downstream stringent bioinformatics analysis [23]. However, these approaches introduce potential sequencing biases related to 3′ de-phosphorylation and PCR amplification steps [24, 25], are time-consuming and computationally expensive.

Here, we present a one-day long and PCR-free library preparation strategy, called circAID-p-seq (CIRCular Amplification and IDentification of short 3′ Phosphate RNA SEQuences). CircAID-p-seq is uniquely characterized by the selection of 3′P/2’-3’cP terminated RNA fragments and cDNA synthesis by rolling circular amplification (RT-RCA) [26, 27], which does not require PCR amplification. In fact, the resulting cDNA is a long (> 200 nt), single-stranded molecule bearing multiple copies of a unique RNA fragment. This cDNA is suitable for direct cDNA sequencing with Oxford Nanopore Technologies (ONT) instruments.

Ribo-seq can be considered an effective case study to test circAID-p-seq, since it requires the sequencing of 25-35 nt-long ribosome-protected fragments (RPFs) produced, *inter alia*, by the RNase I or the Micrococcal nuclease. To evaluate the performance of our new library construction method, we compared our results to standard library preparation and sequencing approaches [28, 29]. To our knowledge, this is an original and unique method for 3′-P/cP RNA-seq library preparation, the first selective 3′-P/cP ribosome profiling, and the first Ribo-seq with the Oxford Nanopore Technologies (ONT) platform.

## Results

RNA molecules with 3′P or 2’,3′cP can be accurately analysed only if the phosphate signature is preserved. To sequence 3′ phosphorylated RNAs, we have developed a new approach for library preparation (circAID-p-seq), which proceeds according to the following five steps (Figure 1a): (1) phosphorylation of the RNA 5′ end, (2) ligation of an RNA adaptor molecule to the 3′P or 2’,3′cP RNA ends [30], (3) intramolecular circularization, (4) retro-transcription rolling circular amplification (RT-RCA) [31] to produce a long concatemeric read containing tandem repeats; each repeat comprises one adaptor and one fragment of interest (called insert hereinafter), (5) second strand cDNA synthesis. Once the cDNA is produced, it can be sequenced with the ONT platform, which allows direct sequencing of the long (> 200 nt) cDNA without additional PCR amplification. After sequencing, all copies of the RNA fragment of interest in the concatemeric raw reads can be identified and processed. To this purpose, we developed *circAidMe*, a computational pipeline implementing identification in the concatemeric read of all RNA fragment copies, multiple sequence alignment of the fragments, and generation of a highly accurate consensus sequence of the RNA fragment for further analyses (Figure 1b). To obtain meaningful biological information, consensus sequences can then be mapped to the target genome or transcriptome for downstream analyses. The circAidMe python code is made freely available on GitHub at: https://github.com/ms-gx/CircAidMe.

**Fig. 1.**
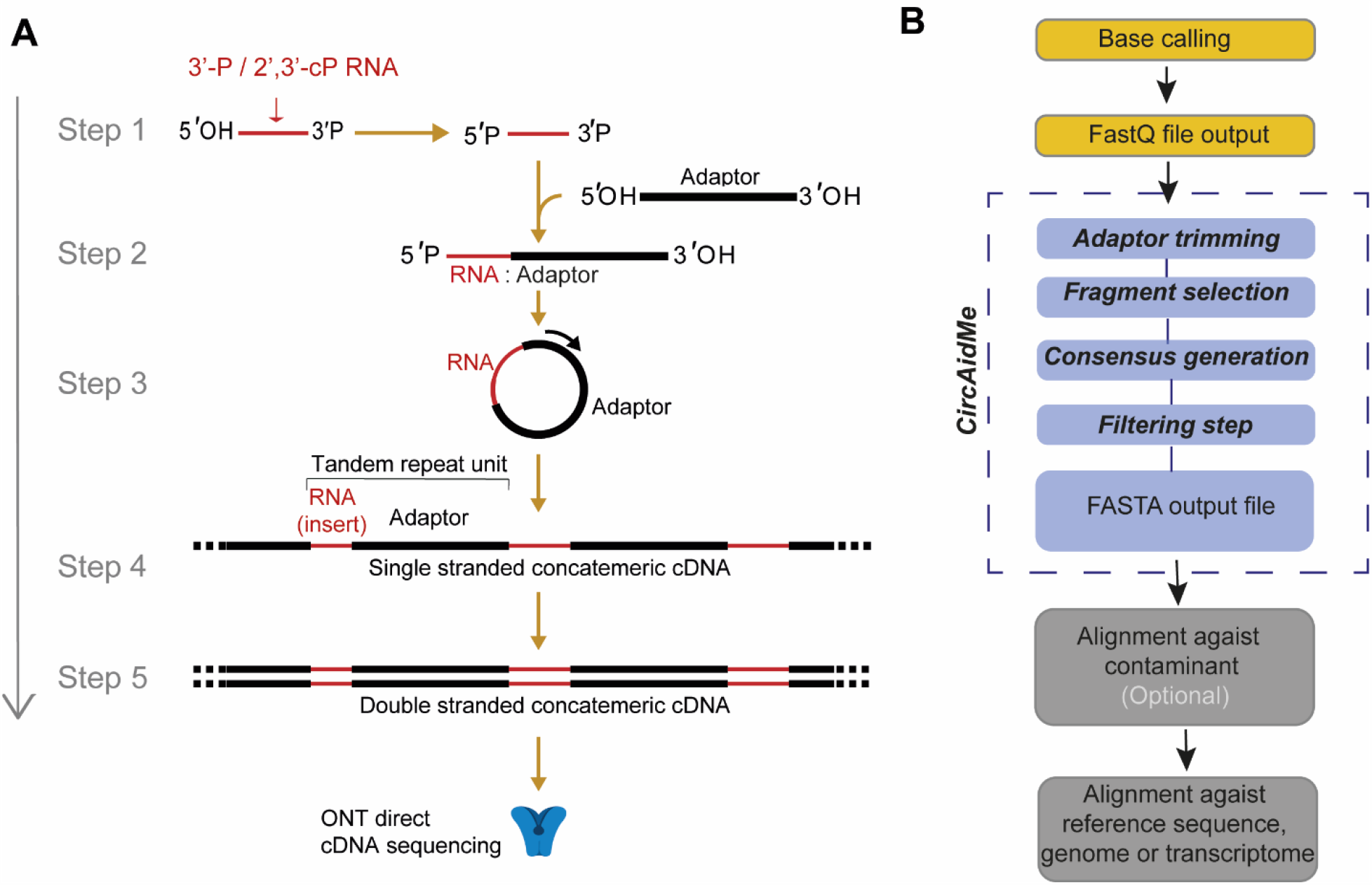
CircAID-p-seq workflow. **a** CircAID-p-seq library preparation: step 1: 5’ phosphorylation of the RNA fragment; step 2: selective capture of the 3’P end through the ligation with the RNA-based adaptor; step 3: circularization; step 4: Reverse Transcription - Rolling Circle Amplification (RT-RCA) - generation of the first strand, and second strand cDNA synthesis; step 5: direct cDNA nanopore sequencing. **b** Bioinformatics pipeline. Bases are called from the ONT output (FAST5 files) and the resulting reads (FASTQ file) are processed with circAidMe, which identifies copies of the RNA fragments and calculates their consensus sequence. Data are filtered of contaminants (e.g., rRNAs, tRNAs, other non-protein-coding transcripts) as needed and aligned against the reference sequences (e.g., genome or transcriptome).

To optimize our strategy, we utilized a 30 nt long synthetic RNA molecule bearing a 3′P group, called RNA30-3′P. The phosphorylation of the 5′ terminus was carried out by the T4 polynucleotide kinase (PNK 3′-minus), a step required to block self-ligation and obtain 5′P-RNA30-3′P RNA species (Figure 1a). The ligation of the 3′P terminus to an adaptor (called ADR hereinafter, with 3′- and 5′-hydroxyl groups) was performed using a 3′P ligase (Supplementary Figure S1A), producing an RNA:adaptor molecule. Intramolecular circularization of the RNA:adaptor was performed with a T4 RNA ligase to form circular RNAs. To confirm the effectiveness of the reaction and to remove remaining single stranded RNAs, the reaction mix was treated with RNase R. This exoribonuclease digests all linear RNA molecules, while preserving circular RNAs [32] (Supplementary Figure S1A). To synthesise the first cDNA strand and generate long single-stranded cDNA molecules carrying multiple copies of the insert, we performed a RT-RCA. To confirm that the multimeric cDNAs had been successfully synthesised, the product was amplified by PCR (Supplementary Figure S1B). Next, we used a Taq polymerase with 5′-3′ exonuclease activity and a primer complementary to the adaptor used in the first ligation, to synthesise the second cDNA strand. The efficiency of second strand synthesis was confirmed by the resistance of cDNA to S1 nuclease digestion, an enzyme that acts on single stranded DNA oligonucleotides but not on double stranded cDNA (Supplementary Figure S2). The library was then sequenced with a benchtop Oxford Nanopore sequencer (MinION). After base calling, the output was analysed by CircAidMe to identify the inserts and generate the consensus sequence (see Materials and Methods and Figure 1b).

To ensure a robust consensus sequence, a large number of inserts in the concatemer reads is desirable. Thus, we assessed the effect of the ADR length and sequence on the number of repeats generated. We used three adaptors, 20 nt (ADR20), 60 nt (ADR60) or 110 nt (ADR110) long and sequenced the same 30 nt synthetic RNA fragment (Figure 2a) (adaptor sequences are listed in Additional file 1). The read length distributions showed a major peak at about 550 nt for all samples, suggesting that ADR20 generated more tandem repeats than ADR60 or ADR110 (Figure 2a and 2b). Then, we investigated the impact of the number of repeats on the accuracy (i.e. the percentage of the read not altered by mismatches or indels, see Materials and Methods) of the consensus sequence. As expected, the high number of repeats obtained with ADR20 results in a more accurate consensus (Figure 2c) and in a narrower distribution of fragments length (Figure 2d). In line with these observations, to maximize the accuracy of the consensus, we used short ADRs for all further experiments.

**Fig. 2.**
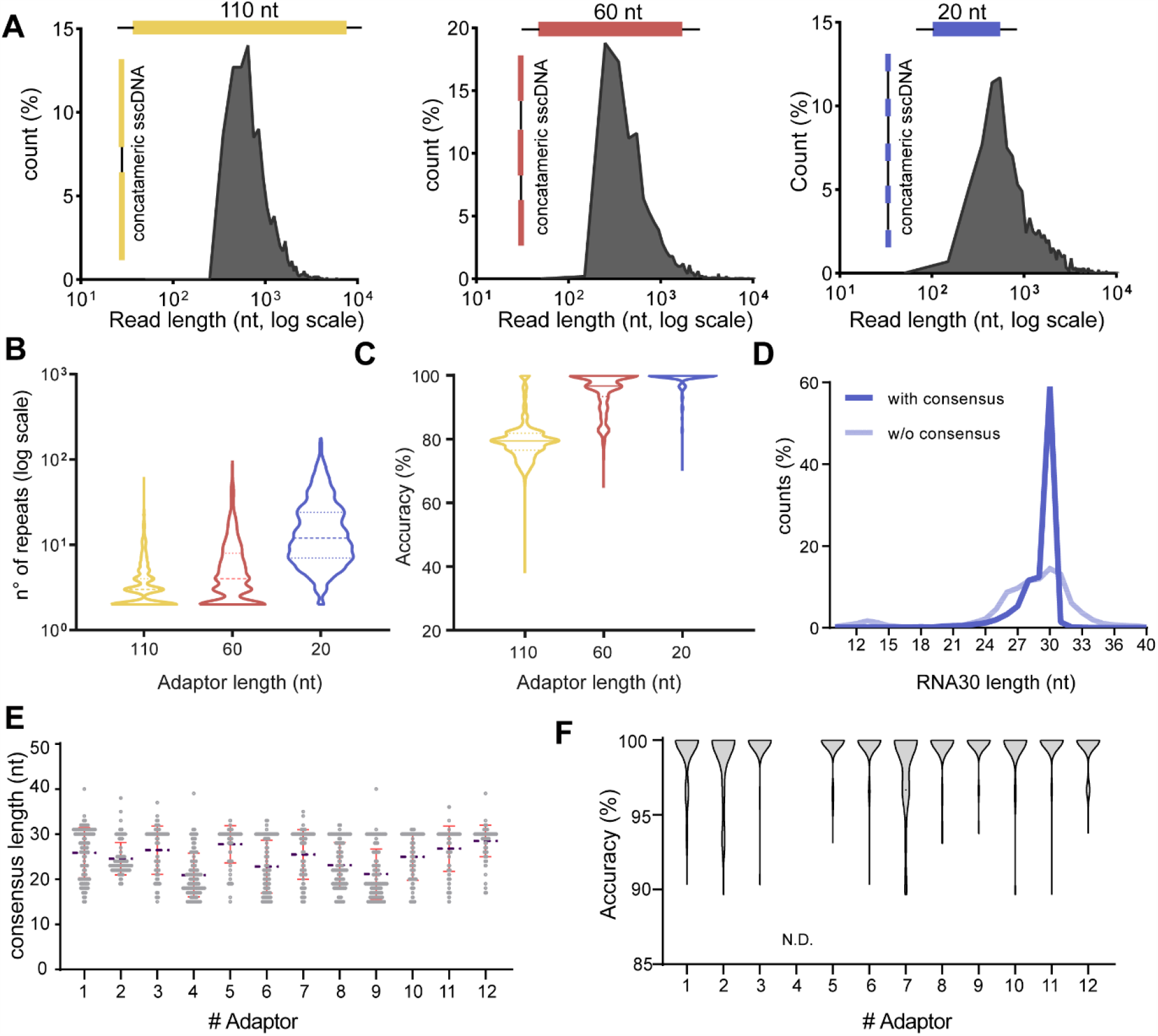
CircAID-p-seq workflow optimization. **a** Three different adaptors (ADR100, yellow; ADR60, red and ADR20, blue) were used to perform CircAID-p-seq on a 30 nt long synthetic fragment phosphorylated at the 3′-end. The raw reads length distribution (nucleotides, log scale) is reported as a function of the relative abundance (%) over total reads. **b** Violin plot showing the effect of the adaptor lengths on the number of fragment repetitions obtained after RT-RCA (dashed line, median). **c** Accuracy of the consensus sequence for the three adaptors (solid line, median). **d** Analysis of the 30RNA 3’-P fragments length distribution before (transparent blue) and after (solid blue) consensus generation, using ADR20 as adaptor. **e** Length distribution of the consensus sequences obtained from 12 different adaptors used to sequence a 30 nt long synthetic insert. (Blue broken line: mean. Red lines: standard deviation above and below mean). **f** Violin plot showing the consensus accuracy of a 30 nt long synthetic fragment obtained with each of the 12 adaptors tested. N.D., not detected, because it did not pass the quality filters for accuracy measurement.

Next, we focused our attention on optimizing the sequence composition of ADRs. We designed twelve 24 nt long ADR oligos (Additional file 1), predicted to have minimal secondary structure. We pooled them at equimolar ratio for capturing and sequencing a 30 nt synthetic fragment (RNA30-3′P). After sequencing, we evaluated the length and the quality of the consensus sequence of the insert, as well as the relative abundance of each ADR. Almost all adaptors achieved a correct consensus length (Figure 2e) and a high accuracy (> 95%) of the insert (Figure 2f) with median of five or more repeats per read (Supplementary Figure 3A). However, reads obtained with some adaptors were more represented than others (Supplementary Figure 3B), suggesting that some of them displayed a higher probability to form RNA:ADR products. Since ADR12 combined good accuracy with a relatively high read abundance, we used this adaptor in all further experiments.

To investigate whether circAID-p-seq can capture quantitative variations in RNA abundance, we sequenced a mixture of three different synthetic RNA fragments (RNA-A, RNA-G and RNA-M) at different molar concentrations. For quantitative analysis we took into consideration only reverse and “hairpin” reads (Supplementary Figure 4A and 4B and methods). Our results showed that there is a good agreement (R^2^ > 0.98) between the amount of input and the number of consensus sequences obtained for each insert (Supplementary figure 4C and 4D). Overall, our results provide evidence that circAID-p-seq (i) can selectively incorporate a mixture of short synthetic RNA molecules bearing a 3′P signature, (ii) is quantitatively accurate, and (iii) is efficiently applicable to the ONT sequencing platform.

### Ribosome profiling with circAID-p-seq

To further confirm the selectivity and efficiency of our library preparation method in complex biological samples, we chose the framework of Ribo-seq, the sequencing of short RNA fragments protected by ribosomes from nuclease digestion. Ribo-seq provides positional information of ribosomes on transcripts as well as an indication of the RNAs engaged in translation. This experimental setup is intrinsically suitable for our purpose because the endonucleases used (RNase A superfamily, e.g. RNase I) are known to generate 3′P ribosome protected fragments (RPF), with a length between 25 nts and 35 nts. Therefore, with circAID-p-seq we can selectively and directly capture only the fragments digested by the nuclease, without the need for additional de-phosphorylation steps and without the risk of sequencing RNAs fragments not generated by the endonuclease.

Since there are currently no available technologies for selecting and sequencing 3′P RNA fragments, we compared our library preparation method with two established protocols for RPF sequencing: 1) the ligation-free sequencing protocol based on a strand switching approach [28] and 2) the classical protocol for ribosome profiling [33]. Both methods are based on the removal of the 3′P prior to library preparation and on Illumina (ILMN) platform. We used HEK293T cells to compare circAID-p-seq to a commercial switching approach (SMARTer smRNA-seq, Clontech) and mouse liver samples for the standard established protocol of ribosome profiling (Figure 3a). To determine whether circAID-p-seq can uncover changes in the localization of ribosomes along transcripts, we compared ribosome footprint distribution in HEK293T cells untreated (H-) and treated (H+) with Harringtonine, a drug known to stall ribosomes at the start codon [34]. We obtained 2 - 5 million raw concatemeric reads per condition by using the CircAID-p-seq libraries, and 47 - 100 million raw reads using Illumina sequencing. The length distribution of the consensus sequences, generated by CircAidMe and representing putative RPFs, peaked at about 33 nts for both HEK293T and liver samples (Figure 3b), in accordance with Illumina data and the length of RNA fragments covered by ribosomes [35]. Even if the two platforms have different sequencing depths, we wanted to better understand the concordances among circAID-p-seq and the ILMN-based methods in term of RPFs coverage, GC content, number and biotype of genes identified. We observed a high correlation in RPFs coverage between circAID-p-seq and the two ILMN sequencing methodologies (Spearman’s R = 0.84 in HEK293T and R = 0.90 in mouse liver samples) (Supplementary Figure S5A and S5B).

**Fig. 3.**
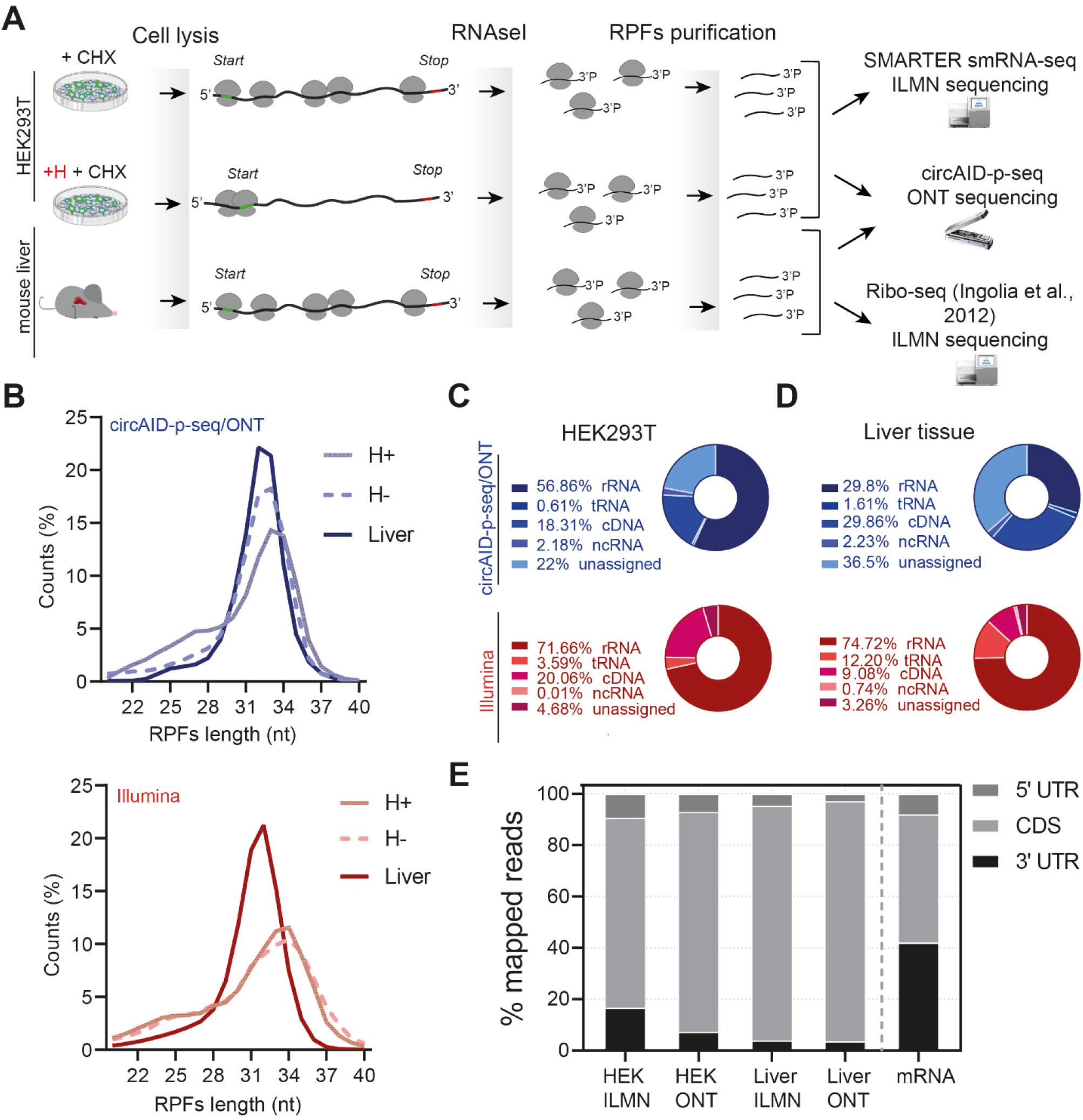
Ribosome Profiling analysis comparison: circAID-p-seq (ONT) vs Ribo-seq (Illumina). **a** Schematic representation of ribosome profiling experiments using HEK293T cells and mouse liver tissue. **b** Ribosome protected fragments (RPFs) length distribution obtained from cells treated (H+) or not (H-) treated with Harringtonine and from liver tissues using circAID-p-seq library prep (top, blue) and the two Illumina Ribo-seq methods (bottom, red). **c** Pie chart representing the percentage of reads mapping on coding and non-coding RNAs in HEK293T and **(d)** mouse liver tissue. **e** Left, percentage of P-sites mapping to the 5’ UTR, coding sequence (CDS), 5’ UTR and 3’ UTR of mRNAs from ONT/circAID-p-seq and Illumina/Ribo-seq data. On the right, theoretical length percentage of each mRNA region (mRNA).

A similar correlation was obtained with circAID-p-seq within replicates (Spearman’s R > 0.90) in mouse liver samples (Supplementary Figure S5C and S5D). A current limitation in small RNA sequencing experiments is the over-representation of sequences generated by PCR amplification [36]. As circAID-p-seq is a PCR-free protocol, PCR duplicates are not an issue. The median percentage of GC content in the liver genes was concordant across the sequencing approaches (52% circAID-p-seq/ONT sequencing vs 56% ILMN sequencing, data not shown), suggesting that there were no GC-related biases in the library preparation method. Of note, CircAID-p-seq showed 45% less reads mapping on rRNA than ILMN in mouse liver samples. The lower rRNA contamination suggests that not all rRNA contaminants derive from cleavage by RNAse I nuclease (i.e. they do not have a 3′P). In circAID-p-seq we also observed less tRNA contamination, but a higher percentage of unassigned reads (Figure 3b, 3c) due to a still high ONT sequencing error rate.

To finally confirm that circAID-p-seq can detect ribosome footprints, we investigated the number and percentage of reads mapping to protein coding sequences (cDNA). In circAID-p-seq data, 18% (0.15 million) of reads in HEK293T and 29% (0.6 million) of reads in mouse liver mapped to coding genes. In Illumina data, 20% (22 million) and 9% (5 million) of the reads in HEK293T and mouse liver respectively, aligned to coding genes (Figure 3b, 3c). We identified a total of 9,419 genes in HEK293T/H-(5,002 with > 10 Transcripts Per kilobase Million, TPM) for circAID-p-seq and 16,754 genes (8,820 with > 10 TPM) (Additional file 2) for ILMN. About 55% of ILMN genes were in common with ONT sequencing (Supplementary Figure 6A). In line with this result, in circAID-p-seq mouse liver we identified 61% (4115 with > 10 TPM) of ILMN genes (Supplementary Figure 6B). We determined that ILMN covers 0.27% of coding genes per million reads generated, while circAID-p-seq/ONT covers 2.45% of the coding genes per million reads generated. In other words, more than 4,000 reads are required to detect a gene in ILMN, while only 130 reads are sufficient with circAID-p-seq. Considering only genes with > 10 TPM, ILMN needs more than 8000 reads/gene while circAID-p-seq requires only 185 reads/gene. Comparable performances were obtained in mouse liver. As a result, even if with ONT the sequencing depth is lower, most of the genes detected by circAID-p-seq match ILMN genes with good read coverage (> 10 TMP), drastically reducing the need for deep sequencing. In agreement with this, if we consider all detected genes (> 1 count), we observed that the majority of circAID-p-seq data have a gene coverage higher than 1 TPM (with a median of 10 TPM) for both HEK293T cells and mouse liver. On the contrary, genes identified only by ILMN are less covered (median of 0.5 TPM), meaning that they have a low density of RPFs (Supplementary Figure 6C and 6D). Overall, these results demonstrate that circAID-p-seq is between four and ten times more efficient in term of number of genes per million of reads with respect to the ILMN sequencing.

To further characterize genes identified by the two methodologies, we performed a Gene Ontology (GO) analysis in both HEK293T and mouse liver datasets. We observed similar enriched terms between ILMN and circAID-p-seq/ONT (Supplementary Figure 6D and 6E), confirming that the most representative transcripts in ILMN were also captured by, and were semantically coherent with, circAID-p-seq data.

Sequencing of ribosome footprints provides a precise record of the position of the ribosome along the mRNAs. RPFs are expected to be over-represented within the coding region (CDS) of the mRNAs, where, in contrast to those positioned in 5′UTR and 3′UTR regions, the P-site positions are expected to exhibit trinucleotide periodicity. In line with this, a high percentage of the generated consensus sequences mapped to the coding sequence (CDS) within mRNAs (85.7% in HEK293T cells; and 93.4 % in mouse liver). The percentage of reads mapping to the 5′UTR and 3′UTR was negligible (Figure 3e). The percentage of P-sites in the three possible translation reading frames obtained with circAID-p-seq were in agreement with ILMN data (Figure 4a, 4b and 4c). A comparison between the P-site metaprofiles at single nucleotide resolution showed a clear trinucleotide periodicity in all sequencing approaches and samples (Figure 4a, 4b and 4c). Treatment with Harringtonine in HEK293T cells showed a relative increase in the signal at the start codon and a decrease along the CDS (Figure 4b) in both circAID-p-seq and ILMN, confirming the robustness of circAID-p-seq in detecting positional information of ribosomes. Metaprofiles in mouse liver obtained using only transcripts detected in both library preparation approaches (n = 4115; > 10 TPMs, Supplementary Figure S7) did not show significant differences between circAID-p-seq and the standard method.

**Fig 4.**
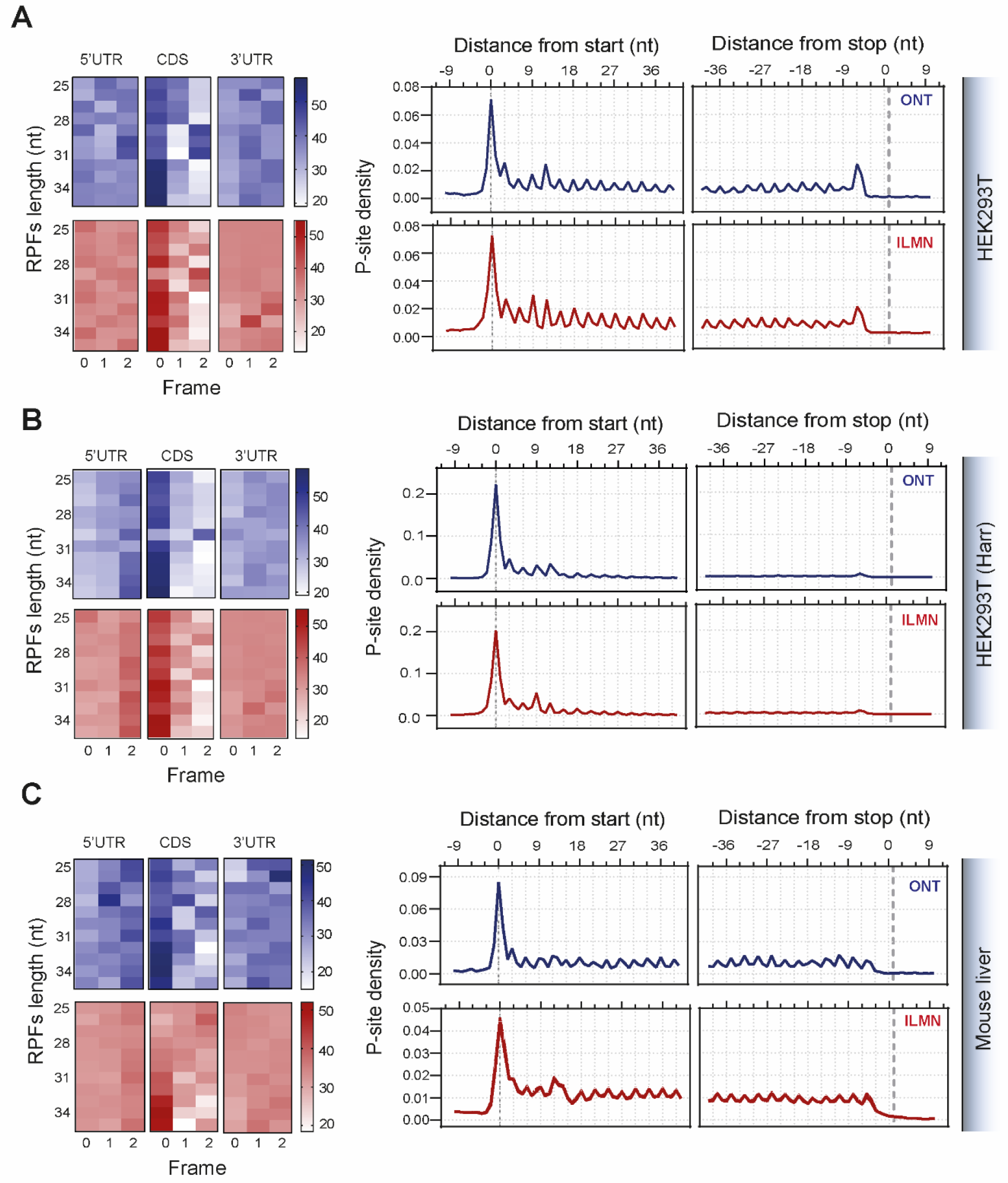
Ribosome footprint data analysis. Percentage of P-sites corresponding to the three possible reading frames (left) along the 5’ UTR, CDS, and 3’ UTR, stratified for read length and metaprofiles (right) showing the density of P-sites around translation initiation site and translation termination site for Riboseq (red) and circAID-p-seq (blue), in HEK293T not treated with Harringtonine (**a**), HEK293T treated with Harringtonine (**b**) and liver tissues (**c**). For liver tissues in **c** data are mean ± s.e.m (shadowed line) of n = 3 biologically independent samples.

Taken together, these results confirm that circAID-p-seq allows selection and sequencing of actual ribosome footprints by including 3′P RNA fragments. Hence, circAID-p-seq retrieves important information to study the positional landscape of ribosomes along transcripts. More specifically, circAID-p-seq, coupled with CircAidMe, generates ribosome profiling data consistent with existing methods; with the added advantages of a higher efficiency and no PCR amplification biases. Remarkably, CircAID-p-seq requires at least 10 times less raw reads than classical ribosome profiling protocols and sequencing, confirming that circAID-p-seq (i) is suitable for ribosome profiling experiments, and (ii) is effective for phospho-RNA-sequencing.

## Discussion

Here, we introduced circAID-p-seq, a new approach for library preparation and nanopore sequencing of 2′,3′ cyclic phosphate- and 3′ phospho-terminated RNA fragments. When combined with the bespoke computational pipeline, CircAidMe, we obtained > 99.5% average accuracy in retrieving the sequence of short synthetic RNAs fragments.

We applied circAID-p-seq to ribosome profiling experiments, which are based on the generation of RNA fragments with 3′-phosphates due to endonuclease digestion. We benchmarked our method against commonly used Ribo-seq library preparation protocols that require a dephosphorylation of the 3′ end. The 3′ dephosphorylation could decrease the level of specificity, resulting in noisier sequencing and inclusion of any short RNA molecule endowed with -OH groups at the 3′ terminus. Moreover, existing approaches are based on PCR steps that can introduce amplification related biases and PCR duplicate issues. With circAID-p-seq, no PCR step is required and after RPF purification no gel extraction steps are needed. Coupled to ONT sequencing, circAID-p-seq produces higher-quality information than approaches based on Illumina sequencing in terms of (i) number of genes per million of reads and (ii) localization of positional data of the ribosomes along transcripts. In addition, circAID-p-seq/ONT sequencing requires 10 to 30 times less absolute number of raw reads than classical ribosome profiling coupled with Illumina sequencing to achieve comparable coverage of the translatome. In fact, CircAID-p-seq showed a higher efficiency in the coverage of the coding genome compared with standard library preparation. Furthermore, circAID-p-seq showed a lower rRNA contamination, at least in our experiments. These features make the circAID-p-seq library generation extremely useful for genome-wide analysis.

If required, an increase in the sequencing depth can be achieved with other ONT sequencing platforms, such as PromethION or GridION. In terms of experimental time, circAID-p-seq combined with CircAidMe allows fast ribosome profiling experiments from sequencing to data analysis (in the region of 24-48 hours). Of note, circAID-p-seq sequencing was performed with the portable and low throughput MinION (ONT) sequencer, affording lower instrumental costs compared to ILMN sequencing (here performed on HiSeq2500 and NovaSeq6000). Another advantage of circAID-p-seq/ONT sequencing is that if samples are not performing well during sequencing, the run can be stopped and the flow cell re-used, a relevant feature in expensive ribosome profiling experiments (Table 1). The main constraints identified for circAID-p-seq are when analysing low abundance transcripts or detecting translation events only marginally represented in Ribo-seq data, such as the translation of upstream open reading frames (uORFs) or ribosome readthrough events [37, 38]. Future developments of circAID-p-seq need to address the option of multiplexing the workflows and of reducing the required amount of input material, currently established in more than 3 picomoles of phosphorylated RNA fragments.

**Table 1.** Summary table of circAID-p-seq/ONT advantages. (*a*: including 24 hours needed for size selection and PAGE purification of RPFs. *b:* with Hornstein N et al. 2016, *c:* with Ingolia et al., 2017)

| Parameter | circAID-p-seq/ONT<br>(MinION) | Illumina (Hiseq2500 -<br>NovaSeq 6000) |
| --- | --- | --- |
| Sequencing Output (million reads) | 2 - 5 | 60 - 100 |
| Time<sup>a</sup> | 39 h | 30 h<sup>b</sup> – 7 days<sup>c</sup> |
| Capital/instrumental cost | low | high |
| Input material (RPFs) | > 50 ng | > 1 ng<sup>b</sup> or > 10 ng<sup>c</sup> |
| Real Time sequencing | Yes | No |

In addition to ribosome profiling, many other RNA footprinting techniques may take advantage of this method, like those employing endoribonucleases and generating 3’P termini with the aim of characterizing RNA-protein interactions, large RNA-protein complexes [40], and/or the interaction of small molecules with RNA [41]. More importantly, 3′P phosphorylated RNAs are hallmarks of biological processes and can be generated in living cells by toxins [5], ribozymes [4], endonucleases [2, 18], the tRNA splicing endonuclease [39], the Ire1 [40], the RNase T2 [20], the RNase L [41] and some CRISPR-associated (Cas) proteins [42]. Endogenous 3′P/2’-3’cP terminated RNA fragments are involved in diverse biological processes, such as RNA metabolism [7], rRNA and tRNA biogenesis [8], mRNA splicing [9], unfolding protein response [10] and stress granules production [11]. Phosphorylated RNA fragments are also dysregulated in disease conditions, such as cancer [43], viral infection [44] and Amyotrophic Lateral Sclerosis [45] pointing to 3′P/2’-3’cP RNAs as likely and largely unexplored signatures of disease [46].

In conclusion, the combination of circAID-p-seq with ONT allows single molecule, low-cost, fast and easy detection of biologically relevant 3′-P/cP RNA species with a portable ONT device. CircAID-p-seq is the first phospho-RNA-sequencing library preparation method successfully tested in ribosome profiling experiments and could be used in the near future to better uncover the biological role of 3′-phospho RNA molecules, a hidden transcriptomic layer in many genome-wide profiles.

## Materials and Methods

### Synthetic oligonucleotides

Custom adaptors, synthetic RNA fragments and DNA primers were synthesized by Integrated DNA Technologies (Coralville, IA). Synthetic RNA fragments consisted of 30-mer oligonucleotides with 5′ OH and 3′P termini. Custom adaptors consisted of RNA molecules with different length and no modification at 5′ and 3′ ends. Adaptors were carefully designed to minimize their secondary structure, using a combination of RNAFold and OligoEvaluator folding tools. All the sequences are listed in Additional file 1.

### circAID-p-seq synthetic library preparation

#### 5′ Phosphorilation and adaptor ligation

Synthetic RNA fragments bearing a 3′P were subjected to 5′ phosphorylation with T4 PNK 3′ minus (NEB, cat n° M0236S), according to manufacturer’s instructions. RNA fragments were purified from the reaction using a RNA Clean & Concentrator(tm)-5 column (Zymo Research, Cat. n° R1013). The resulting RNA fragments were ligated to different adaptors (listed in Supplementary Table 1) via 3′P ligase, according to the following conditions: 30 pmol of RNA fragment, 10 pmol of adaptor, 15 pmol 3′P ligase, 1X 3′P ligase buffer, 100 µM GTP, 1 mM MnCl2 in a final volume of 10 µL. The reaction was incubated 2h at 37°C and then loaded on a 15% acrylamide/8M urea precast gel (Invitrogen, cat n° EC6885BOX). The ligated RNA was purified through gel extraction, as described in the gel analysis section below. To evaluate and optimize the circAID-p-seq method, different combinations of adaptors and synthetic RNA fragments were employed. For testing the effect of different adaptor lengths on RT-RCA and number of repetition, ADR-110, ADR-60, ADR-20 were used for ligation with 30RNA-3’P fragment. Subsequently, to identify the best 24 nt long adaptor sequence, an equimolar pool of 12 oligos was created and used in the first ligation step with 30RNA3’P fragment. For quantitative analysis, three different types of RNA fragments (RNA30-G, RNA30-M, RNA30-A) were combined at the ratio of 1:10:100. In this case, for the circAID-p-seq library preparation, ADR-12 was used.

#### Circularization and RNase R treatment

The circularization of the adaptor-ligated RNA (RNA:adaptor) was carried out at 25 °C for 2h, in a total volume of 20 µL containing 10 U of T4 RNA Ligase 1 (NEB, cat n° M0204L), 1X T4 RNA ligase buffer, 20% PEG8000, 50 µM ATP. The reaction was then incubated at 37 °C for 1 h with 20 U of RNase R (Lucigen, cat n°RNR07250), to remove all the undesired products (i.e linear RNA or concatemer product). Circular RNA was purified through RNA Clean & Concentrator(tm)-5 column (Zymo research, Cat. n° R1013) and quantified using Qubit(tm) RNA HS Assay Kit (Thermo Fisher, Catalog number: Q32852).

#### Reverse Transcription - Rolling Circle Amplification (RT-RCA) and Second Strand synthesis

RT-RCA was performed in 20 µL with Maxima H Minus Reverse Transcriptase (Thermo Fisher, cat n°EP0752) under the following conditions: 50 ng of circular RNA, 200 U of Reverse Transcriptase, 1X RT buffer, 0.5 mM dNTPs, 50 pmol Primer R, 10 % glycerol. The reaction was carried out at 42°C for 4 hours, then stopped by incubation at 70°C for 10 min. After cDNA synthesis, circular RNA template was hydrolyzed by adding 0.1 N NaOH for 10 min at 70°C. The second strand cDNA was generated by performing one PCR cycle using Platinum II Hot start Taq Polymerase (Thermo Fisher, cat n°14966001). The reaction included 20 μL*single-strand* cDNA, 1x Buffer, 0.2 mM dNTPs, 2 mM MgCl_2_, 2 U Taq Polymerase, 50 pmol primer F in a total volume of 50 μL and subjected to the following program: initial denaturation at 94°C, one cycle of 94°C for 30 sec, 60°C for 30 sec and 68°C for 2 minutes. Double strand cDNA was purified using AMPure XP beads (Agencourt, cat n°A63881) according to manufacturer’s instructions. Validation of second strand synthesis was performed by Nuclease S1 digestion (Thermo Fisher, cat n°EN0321) according to manufacturer’s instructions.

#### Nanopore sequencing

Purified double-strand cDNA was prepared for nanopore sequencing. Briefly, cDNA was subjected to end repair and dA-tailing reaction using NEBNext End repair/dA-tailing module (NEB, cat. n°E7546S) following the manufacturer’s instruction and incubated for 5 min at 20°C and then 5 min at 65°C. The reaction mix was purified with AMPure XP beads (Agencourt). ONT Adaptor mix was added according to the direct cDNA sequencing kit protocol (SQK-DCS109, ONT), then loaded on a R9.4 flow cell and sequenced on the MinION sequencing device.

### Gel analysis

The samples were mixed 1:1 with gel loading II (Thermo Fischer Scientific, Cat n°AM8547), denatured at 70°C for 90 secs before being loaded into the gel and run at 200V. Gels were then stained with Sybr™ Gold (Invitrogen, cat n°S11494) and scanned using Chemidoc (GE Healthcare, Piscataway, NJ). Gel images were analyzed using ImageLab (Biorad). When required, bands were isolated from the gel, crushed and soaked overnight in Buffer I (Immagina Biotechnology, cat. n°RL001-10) at room temperature with constant rotation. The aqueous phase was filtered with Millipore ultrafree MC tubes and then precipitated with isopropanol (Sigma, cat. n° I9516) at - 80°C for 2 hours or overnight, then centrifuged for 30 min at 1200g, 4°C. The pellet was washed once with 70% ethanol, centrifuged at 12000 g for 5 min at 4°C and air-dried before further processing.

#### Cell culturing and treatment

HEK293 cells were seeded at 1.5×10^6^ cells/dish and kept in culture until reaching 80% of confluence. Cells were then treated with or without harringtonine (2 µg/mL) for 3 min, followed by CHX (10 μg/mL, SIGMA cat. n° 01810) for 5 min at 37°C. Cell lysates were obtained using a hypotonic lysis buffer (IMMAGINA Biotechnology, cat. n°RL001-1). RNA absorbance at 260 nm was measured by Nanodrop ND-1000 UV-VIS Spectrophotometer and the lysate diluted to a total of 1.7 a.u. (A260 nm) with W-buffer (IMMAGINA Biotechnology, cat. n°RL001-4). RPFs were generated by treating the diluted lysate with 12.7 U of RNase I (Ambion, cat. n°AM2295) at room temperature for 45 min (as described in Clamer et al., 2018). RNase I digestion was stopped by adding 10U of Superase Inhibitor (Thermo Scientific, cat. n° AM2696) for 10 min on ice. Young adult wild-type FVB mice were obtained from breeding stocks at the University of Edinburgh. All procedures were performed under licensed authority from the UK Home Office (PPL P92BB9F93). Liver tissues were dissected immediately following sacrifice and pulverized under liquid nitrogen using a pestle and a mortar and the lysates obtained according to previous protocols [47]. Lysates were treated with RNAse I and the 80S with RPFs were isolated using sucrose gradient separation according to previous protocols [48] (Lauria et al., 2020). The RPF were purified using acidic phenol/chloroform extraction. The RPF (25-35 nt) were obtained after purification in denaturing 15% UREA-PAGE. The RPF were divided into two aliquots. One was used for library preparation [29, 48]. The other aliquot was used for circAID-p-seq. All experiments were performed in biological triplicate.

#### Sucrose cushioning

After digestion, lysates were loaded on top of 900 µL of a 30 % sucrose cushion (30 mM Tris-HCl pH 7.5, 100 mM NaCl, 10 mM MgCl_2_, 1M sucrose in nuclease-free water) supplemented with 20 μg/mL of CHX. Samples were ultracentrifuged at 95,000 rpm at 4°C for 2h using a TLA_100.2_ rotor (Beckman). The pellets were resuspended in 200 µL of W-Buffer and treated with 1% SDS (Sigma cat. n° 05030) and 0.1 mg Proteinase K (Euroclone, cat. n° EMR022001) at 37°C for 75 min. Total RNA was extracted by acid-phenol:chloroform, pH 4.5 (Ambion, cat n° AM9722). RNA was precipitated with isopropanol, air-dried, resuspended in nuclease-free water and analyzed on 15% acrylamide/8M urea precast gel. RPFs were size-selected (corresponding to 25-35 nt bands) and extracted from gel (see above). Before starting with library preparation, isolated and purified RPFs were quantified using the Qubit™ miRNA Assay Kit (Thermo Fisher, Catalog number: Q32881).

#### Ribo-seq sample and library preparation

Libraries from RPFs isolated from HEK293T cells were prepared using the SMARTer® smRNA-Seq Kit for Illumina (Takara, Cat. No. 635029) and sequenced with 50 cycles single-read on an Illumina NovaSeq 6000 sequencer. HEK293T RPFs were extracted from independent biological replicates for circAID-p-seq/ONT and SMARTer® smRNA-Seq /Illumina sequencing. Lists of the counts per gene are reported in Additional file 2. Mouse liver RPFs were extracted from three biological replicates. For each replicate, the same pool of PAGE purified RPFs of each triplicate were used for parallel circAID-p-seq/ONT and Illumina sequencing. Illumina libraries were prepared according to Ingolia et al., 2012 and were sequenced with 50 cycles single-read on an Illumina HiSeq2500 sequencer. List of the counts per gene reported are in Additional file 3.

For circAID-p-seq, upon 5′ phosphorylation of RPFs, library preparation was performed starting from 10 pmol of ADR12, following the protocol as described above. The final libraries were loaded on a R9.4 flow cell and sequenced for 20 h with a MinION sequencer.

#### circAID-p-seq data analysis

To identify the RPFs (fragments) contained in each read we developed a custom pipeline (written in Python 3): CircAidMe. The FASTQ files obtained by Guppy 3.6.1 (available from ONT via https://community.nanoporetech.com) base calling are processed to identify from each read the consensus of the repeat sequence from every fragment meeting certain selection criteria. These criteria are user-defined and include filtering by fragment length (for our libraries we selected fragment lengths between 15 and 40 nt) and minimum number of fragment copies required to calculate a consensus sequence. CircAidMe performs the following steps: First, fused reads are detected by 1) searching for remaining ONT sequencing adaptors in a read and 2) detecting orientation patterns of the circAID-p-seq adaptors, which indicate a fused read. If a fused read is encountered, it is split at the appropriate position. Then, for every (split or non-split) read circAID-p-seq adaptors are detected. The fragment inserts flanked by the adaptors are then extracted and a first multiple sequence alignment (MSA) is performed with MUSCLE v3.8.1551 [49]. The MSA is examined using esl-alipid (https://github.com/EddyRivasLab/easel/tree/master/miniapps) and low quality inserts are detected and removed. A second MUSCLE run is performed, resulting in the final consensus sequence of the fragment. The detection of ONT- and circAID-p-seq adaptors is performed using SeqAn v2.4 [50]. Additionally, several statistics are collected while executing CircAidMe to evaluate the quality of the circAID-p-seq library, including: number of fragments per read (> 2 for Ribo-seq), number of adaptors detected per read (≥ 4 for Ribo-seq), read length and consensus length (≥ 20 nt for Ribo-seq). These quantities are recorded for each read in a text file in CSV format. Moreover, for each read that was excluded from the final output the reason is given as a tag in the report file. The final retained consensus sequences are collected in a FASTA file for downstream analyses. Reads discarded by CircAidMe are collected in a separate FASTA file. More details about the function of CircAidMe can be found on its GitHub page: https://github.com/ms-gx/CircAidMe. Read accuracy was determined as reported in Volden R. et al., 2018. Briefly, the accuracy is given as a percentage, representing the portion of the consensus sequence not altered by mismatches or indels when considering the alignment to the reference sequence/coding transcriptome. The process is executed for each aligned consensus sequence.

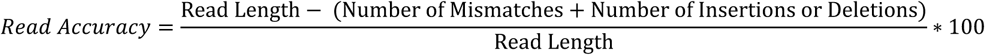

(i) accuracy calculation

For quantitative analysis, we first reasoned that the two strands of the concatemeric cDNA carry complementary information (Supplementary Figure 4A). If this was the case, only one strand of the double stranded cDNA should to be taken into account for an accurate quantitative analysis and in order to not count two times the same fragment. To evaluate if this had occurred, and to then determine which strand is more reliable, we compared the raw read length of both forward and reverse cDNA concatemeric strands based on the orientation of the ADR-12 sequence (Supplementary Figure 4A). We noticed that forward reads, generated during the second strand synthesis, are generally shorter and less abundant than the reverse reads (Supplementary Figure 4B). This effect most probably derives from multiple annealing sites of the primer for second strand synthesis on the concatemeric cDNA. Therefore, since forward-strand synthesis can generate fragmented copy of the reverse strand, it could introduce biases in quantitative experiments. Then, we noticed that a portion of the reads carry forward and reverse strands fused together. This effect is known to be caused by two different molecular reactions: (i) a second strand entering the pore immediately after the first strand without the sequencing device been able to detect two independent molecules (we called this type of reads “fused reads”) [25], or (ii) a hairpin at the 3’-end of the cDNA strand that functions as a primer during second strand synthesis [26] (we called this type of reads “hairpin reads”, Supplementary Figure 4A). We optimized CircAidMe to split fused reads, while leaving hairpin reads untouched. The latter are useful to produce longer reads (Supplementary Figure 4B), hence useful for a better consensus and for quantitative information.

#### Ribosome profiling data analysis

To assess representation of different components of the human transcriptome in the libraries, the consensus sequences generated by CircAidMe were iteratively mapped with Bowtie2 [51] to different classes of human transcribed sequences (as annotated In Gencode v33), including: rRNAs, tRNAs, other non-protein-coding transcripts, and the protein-coding transcriptome. The remaining unmapped sequences were aligned to the human Genome (assembly GRCh38.p13). For ribosome profiling down-stream analyses, consensus sequences were aligned directly to the genome after removal of those mapping to rRNA and tRNA sequences. Alignment files were processed with Samtools version 1.9 [52] and analyzed with the RiboWaltz R package [53], to identify the P-site localization within ribosome footprints and to assess three-periodicity and representation of coverage within coding regions of the mRNAs. As an extra QC step, the Read Accuracy index [54] was calculated for reads of selected libraries with a custom R script. Illumina data from human HEK293T cells were processed with the SMARTer® smRNA-Seq Kit for Illumina (Takara, Cat. Nos. 635029) following guidelines. Briefly, reads were trimmed with Cutadapt version 2.10 [55] by removing the first 3 nucleotides of every read and the 3’ terminal adaptor plus poly-A tail and trimmed reads of length under 15 nucleotides were discarded. Illumina data from mouse liver tissue were processed according to Ingolia et al., 2012, removing the Illumina adaptors and retaining only reads where the adaptors were present (parameters used: -O 15 –e 0.15 -n 3 –m 19). Illumina reads from HEK293T and mouse liver samples were analysed following the same procedure detailed above for circAID-p-seq data, aligning them to the respective classes of transcribed sequences and genomes (Gencode v33 annotation and genome assembly GRCh38.p13 for human; Genecode vM34 annotation and genome assembly GRCm38.p6 for mouse). Read coverages for each protein-coding gene from ONT and Illumina libraries were calculated with HTSeq [56] and normalized as TPM values for comparison between libraries. Only coverages by more than one read were retained in comparisons between ONT and Illumina libraries. PCR duplicate reads were identified with Picard MarkDuplicates version 2.23.1 (Picard Toolkit.” 2019. Broad Institute, GitHub Repository. http://broadinstitute.github.io/picard/; Broad Institute) with default parameters, and removed with Samtools.

## Ethics approval and consent to participate

All animal tissue was obtained under full licensed approval from the UK Home Office following internal (institutional) and external ethical review.

## Availability of data and materials

Source code of CircAidMe along with a pre-compiled package and a description of the software are available on GitHub: https://github.com/ms-gx/CircAidMe.

## Authors’ contributions

Conceptualization: M.C., A.D.P. Bioinformatic analysis: R.B., M.S., L.B., F.L. Implementation of CircAidMe: M.S. Wrote first draft of paper: M.C., A.D.P. Tissue/Data collection: A.D.P., C.F., L.M., M.C., I.S., S.T., T.H.G. Data curation: M.C., PB, A.D.P. Supervised research: M.C. Paper revision: G.V., M.S., A.D.P., M.C., F.L. T.H.G., L.B., S.T. Reviewed and approved text: all authors. The authors read and approved the final manuscript.

## Acknowledgements

We thank Pervasive Edge Srls for the service (reference number: 7/E).

## References

1. Lee C, Harris RA, Wall JK, Mayfield RD, Wilke CO. RNaseIII and T4 Polynucleotide Kinase sequence biases and solutions during RNA-seq library construction. Biol Direct. 2013;8:16. doi:10.1186/1745-6150-8-16.

2. Cuchillo CM, Nogués MV, Raines RT. Bovine Pancreatic Ribonuclease: Fifty Years of the First Enzymatic Reaction Mechanism. Biochemistry. 2011;50:7835–41. doi:10.1021/bi201075b.

3. Grosshans CA, Cech TR. A hammerhead ribozyme allows synthesis of a new form of the Tetrahymena ribozyme homogeneous in length with a 3’ end blocked for transesterification. Nucleic Acids Res. 1991;19:3875–80. doi:10.1093/nar/19.14.3875.

4. Price SR, Ito N, Oubridge C, Avis JM, Nagai K. Crystallization of RNA-protein complexes I. Methods for the large-scale preparation of RNA suitable for crystallographic studies. J Mol Biol. 1995;249:398–408. doi:10.1006/jmbi.1995.0305.

5. Ogawa T, Inoue S, Yajima S, Hidaka M, Masaki H. Sequence-specific recognition of colicin E5, a tRNA-targeting ribonuclease. Nucleic Acids Res. 2006;34:6065–73. doi:10.1093/nar/gkl629.

6. Ogawa T. A Cytotoxic Ribonuclease Targeting Specific Transfer RNA Anticodons. Science (80-). 1999;283:2097–100. doi:10.1126/science.283.5410.2097.

7. Yoshinari S, Liu Y, Gollnick P, Ho CK. Cleavage of 3′-terminal adenosine by archaeal ATP-dependent RNA ligase. Sci Rep. 2017;7:11662. doi:10.1038/s41598-017-11693-0.

8. Filipowicz W, Shatkin AJ. Origin of splice junction phosphate in tRNAs processed by HeLa cell extract. Cell. 1983;32:547–57. doi:10.1016/0092-8674(83)90474-9.

9. Shinya S, Kadokura H, Imagawa Y, Inoue M, Yanagitani K, Kohno K. Reconstitution and characterization of the unconventional splicing of XBP1u mRNA in vitro. Nucleic Acids Res. 2011;39:5245–54. doi:10.1093/nar/gkr132.

10. Maurel M, Chevet E, Tavernier J, Gerlo S. Getting RIDD of RNA: IRE1 in cell fate regulation. Trends Biochem Sci. 2014;39:245–54. doi:10.1016/j.tibs.2014.02.008.

11. Ivanov P, Emara MM, Villen J, Gygi SP, Anderson P. Angiogenin-Induced tRNA Fragments Inhibit Translation Initiation. Mol Cell. 2011;43:613–23. doi:10.1016/j.molcel.2011.06.022.

12. Ramanathan M, Porter DF, Khavari PA. Methods to study RNA–protein interactions. Nat Methods. 2019;16:225–34. doi:10.1038/s41592-019-0330-1.

13. Van Nostrand EL, Freese P, Pratt GA, Wang X, Wei X, Xiao R, et al. A large-scale binding and functional map of human RNA-binding proteins. Nature. 2020;583:711–9. doi:10.1038/s41586-020-2077-3.

14. Silverman IM, Li F, Alexander A, Goff L, Trapnell C, Rinn JL, et al. RNase-mediated protein footprint sequencing reveals protein-binding sites throughout the human transcriptome. Genome Biol. 2014;15:R3. doi:10.1186/gb-2014-15-1-r3.

15. Singh G, Ricci EP, Moore MJ. RIPiT-Seq: A high-throughput approach for footprinting RNA:protein complexes. Methods. 2014;65:320–32. doi:10.1016/j.ymeth.2013.09.013.

16. Nicholson CO, Friedersdorf M, Keene JD. Quantifying RNA binding sites transcriptome-wide using DO-RIP-seq. RNA. 2017;23:32–46. doi:10.1261/rna.058115.116.

17. Maticzka D, Ilik IA, Aktas T, Backofen R, Akhtar A. uvCLAP is a fast and non-radioactive method to identify in vivo targets of RNA-binding proteins. Nat Commun. 2018;9:1142. doi:10.1038/s41467-018-03575-4.

18. Lu L, Li J, Moussaoui M, Boix E. Immune Modulation by Human Secreted RNases at the Extracellular Space. Front Immunol. 2018;9. doi:10.3389/fimmu.2018.01012.

19. Ikehara M, Ohtsuka E, Tokunaga T, Nishikawa S, Uesugi S, Tanaka T, et al. Inquiries into the structure-function relationship of ribonuclease T1 using chemically synthesized coding sequences. Proc Natl Acad Sci. 1986;83:4695–9. doi:10.1073/pnas.83.13.4695.

20. Luhtala N, Parker R. T2 Family ribonucleases: ancient enzymes with diverse roles. Trends Biochem Sci. 2010;35:253–9. doi:10.1016/j.tibs.2010.02.002.

21. Spitale RC, Flynn RA, Zhang QC, Crisalli P, Lee B, Jung J-W, et al. Structural imprints in vivo decode RNA regulatory mechanisms. Nature. 2015;519:486–90. doi:10.1038/nature14263.

22. Honda S, Morichika K, Kirino Y. Selective amplification and sequencing of cyclic phosphate– containing RNAs by the cP-RNA-seq method. Nat Protoc. 2016;11:476–89. doi:10.1038/nprot.2016.025.

23. Giraldez MD, Spengler RM, Etheridge A, Goicochea AJ, Tuck M, Choi SW, et al. Phospho-RNA-seq: a modified small RNA-seq method that reveals circulating mRNA and lncRNA fragments as potential biomarkers in human plasma. EMBO J. 2019;38. doi:10.15252/embj.2019101695.

24. Leshkowitz D, Horn-Saban S, Parmet Y, Feldmesser E. Differences in microRNA detection levels are technology and sequence dependent. RNA. 2013;19:527–38. doi:10.1261/rna.036475.112.

25. Aird D, Ross MG, Chen W-S, Danielsson M, Fennell T, Russ C, et al. Analyzing and minimizing PCR amplification bias in Illumina sequencing libraries. Genome Biol. 2011;12:R18. doi:10.1186/gb-2011-12-2-r18.

26. You X, Vlatkovic I, Babic A, Will T, Epstein I, Tushev G, et al. Neural circular RNAs are derived from synaptic genes and regulated by development and plasticity. Nat Neurosci. 2015;18:603–10. doi:10.1038/nn.3975.

27. Kumar P, Johnston BH, Kazakov SA. miR-ID: A novel, circularization-based platform for detection of microRNAs. RNA. 2011;17:365–80. doi:10.1261/rna.2490111.

28. Hornstein N, Torres D, Das Sharma S, Tang G, Canoll P, Sims PA. Ligation-free ribosome profiling of cell type-specific translation in the brain. Genome Biol. 2016;17:149. doi:10.1186/s13059-016-1005-1.

29. Ingolia NT, Brar GA, Rouskin S, McGeachy AM, Weissman JS. The ribosome profiling strategy for monitoring translation in vivo by deep sequencing of ribosome-protected mRNA fragments. Nat Protoc. 2012;7:1534–50. doi:10.1038/nprot.2012.086.

30. Chakravarty AK, Subbotin R, Chait BT, Shuman S. RNA ligase RtcB splices 3’-phosphate and 5’-OH ends via covalent RtcB-(histidinyl)-GMP and polynucleotide-(3’)pp(5’)G intermediates. Proc Natl Acad Sci. 2012;109:6072–7. doi:10.1073/pnas.1201207109.

31. Acevedo A, Andino R. Library preparation for highly accurate population sequencing of RNA viruses. Nat Protoc. 2014;9:1760–9. doi:10.1038/nprot.2014.118.

32. Suzuki H. Characterization of RNase R-digested cellular RNA source that consists of lariat and circular RNAs from pre-mRNA splicing. Nucleic Acids Res. 2006;34:e63–e63. doi:10.1093/nar/gkl151.

33. Ingolia NT, Ghaemmaghami S, Newman JRS, Weissman JS. Genome-Wide Analysis in Vivo of Translation with Nucleotide Resolution Using Ribosome Profiling. Science (80-). 2009;324:218–23. doi:10.1126/science.1168978.

34. Ingolia NT, Lareau LF, Weissman JS. Ribosome Profiling of Mouse Embryonic Stem Cells Reveals the Complexity and Dynamics of Mammalian Proteomes. Cell. 2011;147:789–802. doi:10.1016/j.cell.2011.10.002.

35. Wu CC-C, Zinshteyn B, Wehner KA, Green R. High-Resolution Ribosome Profiling Defines Discrete Ribosome Elongation States and Translational Regulation during Cellular Stress. Mol Cell. 2019;73:959-970.e5. doi:10.1016/j.molcel.2018.12.009.

36. Fu Y, Wu P-H, Beane T, Zamore PD, Weng Z. Elimination of PCR duplicates in RNA-seq and small RNA-seq using unique molecular identifiers. BMC Genomics. 2018;19:531. doi:10.1186/s12864-018-4933-1.

37. Wangen JR, Green R. Stop codon context influences genome-wide stimulation of termination codon readthrough by aminoglycosides. Elife. 2020;9. doi:10.7554/eLife.52611.

38. Lin Y, May GE, Kready H, Nazzaro L, Mao M, Spealman P, et al. Impacts of uORF codon identity and position on translation regulation. Nucleic Acids Res. 2019;47:9358–67. doi:10.1093/nar/gkz681.

39. Trotta CR, Miao F, Arn EA, Stevens SW, Ho CK, Rauhut R, et al. The Yeast tRNA Splicing Endonuclease: A Tetrameric Enzyme with Two Active Site Subunits Homologous to the Archaeal tRNA Endonucleases. Cell. 1997;89:849–58. doi:10.1016/S0092-8674(00)80270-6.

40. Sidrauski C, Walter P. The Transmembrane Kinase Ire1p Is a Site-Specific Endonuclease That Initiates mRNA Splicing in the Unfolded Protein Response. Cell. 1997;90:1031–9. doi:10.1016/S0092-8674(00)80369-4.

41. Cooper DA, Banerjee S, Chakrabarti A, García-Sastre A, Hesselberth JR, Silverman RH, et al. RNase L Targets Distinct Sites in Influenza A Virus RNAs. J Virol. 2015;89:2764–76. doi:10.1128/JVI.02953-14.

42. Hale CR, Zhao P, Olson S, Duff MO, Graveley BR, Wells L, et al. RNA-Guided RNA Cleavage by a CRISPR RNA-Cas Protein Complex. Cell. 2009;139:945–56. doi:10.1016/j.cell.2009.07.040.

43. Honda S, Loher P, Shigematsu M, Palazzo JP, Suzuki R, Imoto I, et al. Sex hormone-dependent tRNA halves enhance cell proliferation in breast and prostate cancers. Proc Natl Acad Sci. 2015;112:E3816–25. doi:10.1073/pnas.1510077112.

44. Wang Q, Lee I, Ren J, Ajay SS, Lee YS, Bao X. Identification and Functional Characterization of tRNA-derived RNA Fragments (tRFs) in Respiratory Syncytial Virus Infection. Mol Ther. 2013;21:368–79. doi:10.1038/mt.2012.237.

45. Thiyagarajan N, Ferguson R, Subramanian V, Acharya KR. Structural and molecular insights into the mechanism of action of human angiogenin-ALS variants in neurons. Nat Commun. 2012;3:1121. doi:10.1038/ncomms2126.

46. Godoy PM, Bhakta NR, Barczak AJ, Cakmak H, Fisher S, MacKenzie TC, et al. Large Differences in Small RNA Composition Between Human Biofluids. Cell Rep. 2018;25:1346–58. doi:10.1016/j.celrep.2018.10.014.

47. Bernabò P, Tebaldi T, Groen EJN, Lane FM, Perenthaler E, Mattedi F, et al. In Vivo Translatome Profiling in Spinal Muscular Atrophy Reveals a Role for SMN Protein in Ribosome Biology. Cell Rep. 2017;21:953–65. doi:10.1016/j.celrep.2017.10.010.

48. Lauria F, Bernabò P, Tebaldi T, Groen EJN, Perenthaler E, Maniscalco F, et al. SMN-primed ribosomes modulate the translation of transcripts related to spinal muscular atrophy. Nat Cell Biol. 2020;22:1239–51. doi:10.1038/s41556-020-00577-7.

49. Edgar RC. MUSCLE: multiple sequence alignment with high accuracy and high throughput. Nucleic Acids Res. 2004;32:1792–7. doi:10.1093/nar/gkh340.

50. Reinert K, Dadi TH, Ehrhardt M, Hauswedell H, Mehringer S, Rahn R, et al. The SeqAn C++ template library for efficient sequence analysis: A resource for programmers. J Biotechnol. 2017;261:157–68. doi:10.1016/j.jbiotec.2017.07.017.

51. Langmead B, Salzberg SL. Fast gapped-read alignment with Bowtie 2. Nat Methods. 2012;9:357–9. doi:10.1038/nmeth.1923.

52. Li H, Handsaker B, Wysoker A, Fennell T, Ruan J, Homer N, et al. The Sequence Alignment/Map format and SAMtools. Bioinformatics. 2009;25:2078–9. doi:10.1093/bioinformatics/btp352.

53. Lauria F, Tebaldi T, Bernabò P, Groen EJN, Gillingwater TH, Viero G. riboWaltz: Optimization of ribosome P-site positioning in ribosome profiling data. PLOS Comput Biol. 2018;14:e1006169. doi:10.1371/journal.pcbi.1006169.

54. Volden R, Palmer T, Byrne A, Cole C, Schmitz RJ, Green RE, et al. Improving nanopore read accuracy with the R2C2 method enables the sequencing of highly multiplexed full-length single-cell cDNA. Proc Natl Acad Sci. 2018;115:9726–31. doi:10.1073/pnas.1806447115.

55. Martin M. Cutadapt removes adapter sequences from high-throughput sequencing reads. EMBnet.journal. 2011;17:10. doi:10.14806/ej.17.1.200.

56. Anders S, Pyl PT, Huber W. HTSeq--a Python framework to work with high-throughput sequencing data. Bioinformatics. 2015;31:166–9. doi:10.1093/bioinformatics/btu638.

