## Supplementary Data and Figures for "Phospho-RNA sequencing with CircAID-p-seq"

### Supplementary information

### Supplementary figures

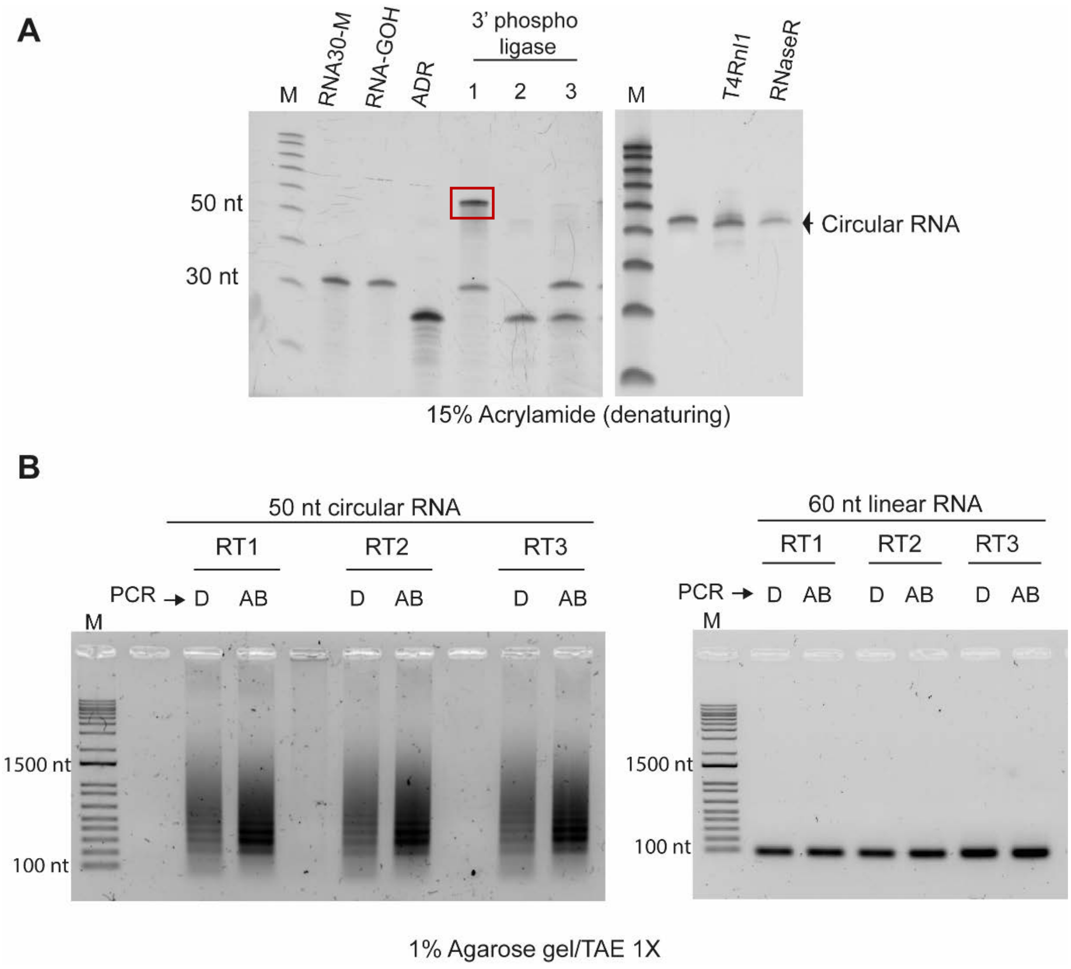


Figure S1. **circAID-p-seq workflow validation. (A)** TBE-urea PAGE gels showing the 3'P-mediated ligation of a synthetic 30 nt long RNA-3'P and a 24 nt long adaptor (ADR12, left panel). Lane labels indicate: RNA30-M, synthetic RNA oligo of 30 nt with a 3'P; RNA-GOH synthetic RNA oligo of 30 nt with a 3'OH; ADR, adaptor; 1, 3'P- ligation reaction between ADR12 and RNA30-M (red box shows the reaction product); 2, 3'P- ligation reaction only with ADR12 without any RNA fragment (negative control); 3, 3'P- ligation reaction between ADR12 and RNA30-GOH. On the right panel, RNA-3'P-ADR product extracted and loaded on the TBE-urea gel again, followed by circularization with T4 Rnl1 and digestion with RNase R. Note that only 1/10 of the total RNA products were loaded for each step to not overload the lane. **(B)** Agarose gels showing PCR products obtained from circular (left panel) and linear (right panel) cDNA, by two different Taq enzymes: D, dreamTaq (Thermo Fisher); AB, Super AB Taq (AB Analitica).

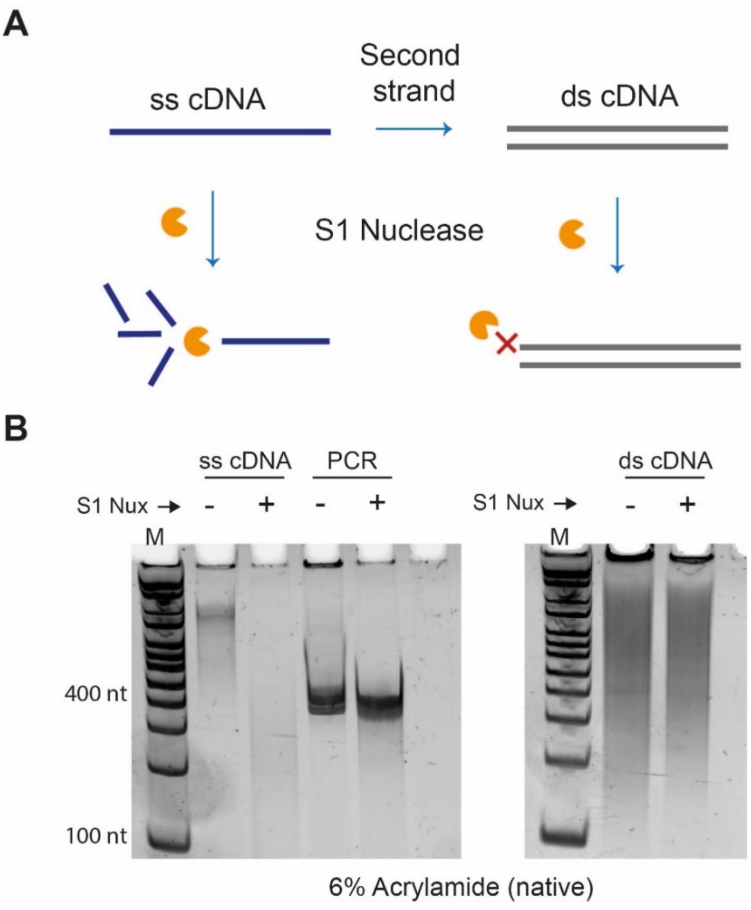


Figure S2. **circAID-p-seq, second strand synthesis validation. (A)** schematic representation: S1 nuclease digests single-stranded cDNA but not double-stranded cDNA. **(B)** TBE-urea PAGE gels showing the effect of S1 nuclease (S1 Nux) treatment (+) compared with no treatment (-) on the single-stranded cDNA and PCR products (left). On the right, the double-stranded cDNA obtained with circAID-p-seq with (+) and without (-) S1 treatment.

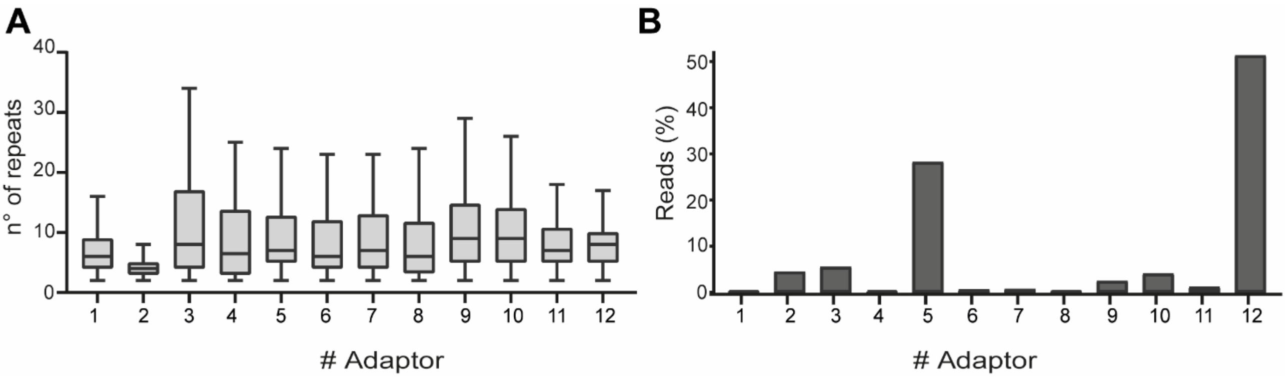


Figure S3. **circAID-p-seq adaptor optimization.** Output from an equimolar pool of 12 different adaptors. **(A)** Number of repeats obtained from each adaptor. **(B)** Number of reads obtained from each adaptor.

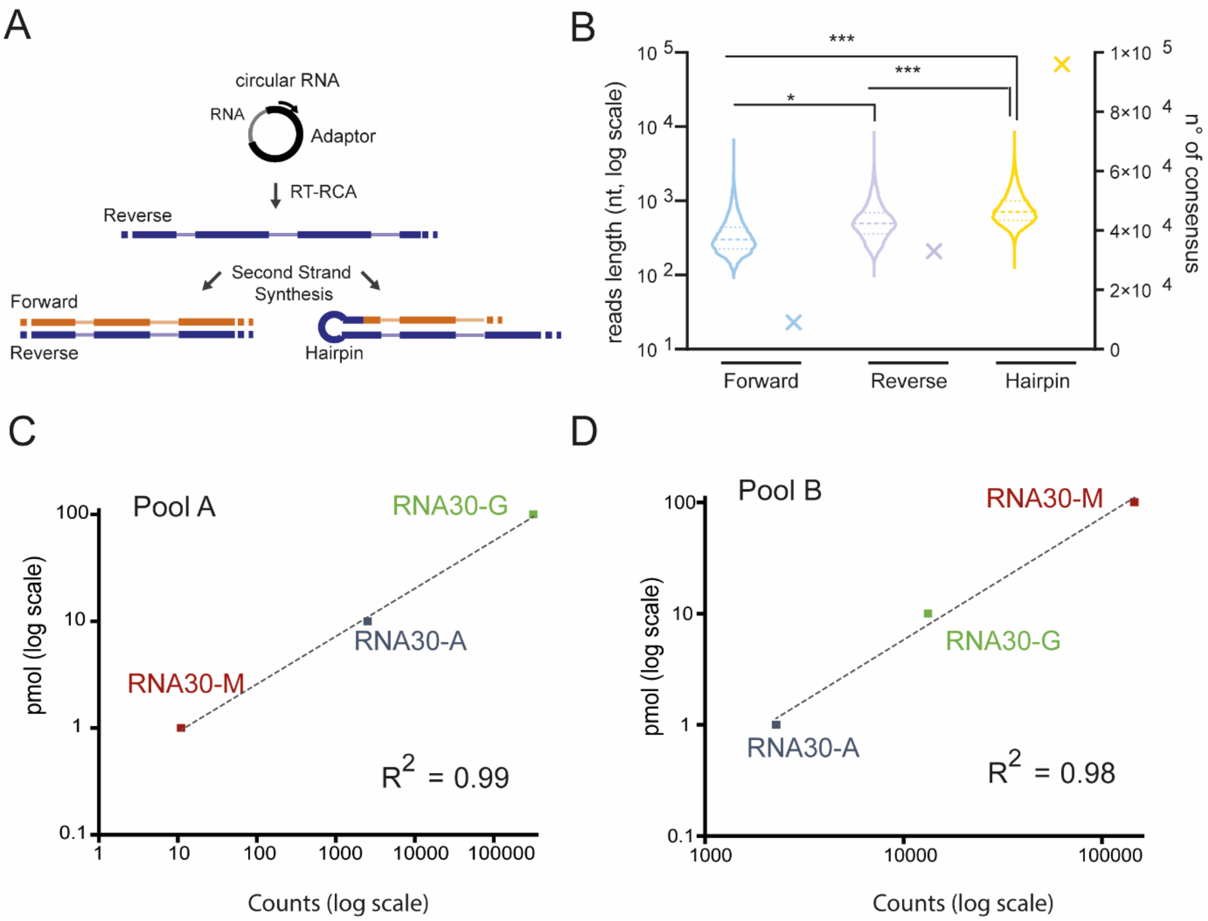


Figure S4. **Quantitative analysis. (A)** Schematic representation of reverse (blue), forward (orange) and hairpin strand, formed during circAID-p-seq steps of RT-RCA and second strand synthesis. **(B)** Violin and dot plot reporting the reads length counts (dashed line: median). The crosses report the number of consensus sequences generated after circAidMe analysis for each strand. Significance was tested with Wilcoxon test P-value (*=P < 0.05; ***= P ≤ 2.792e-13) **(C)** Correlation between read counts and expected abundances (pmol) for Pool A **(**RNA30-M, RNA30-A and RNA30-G, mixed at 1:10:100 picomolar ratio respectively). **(D)** Correlation between read counts and expected abundances (pmol) for Pool B **(**RNA30-A, RNA30-G and RNA30-M), mixed at 1:10:100 picomolar ratio respectively. Squared Pearson correlation coefficients ($R^2$) are reported.

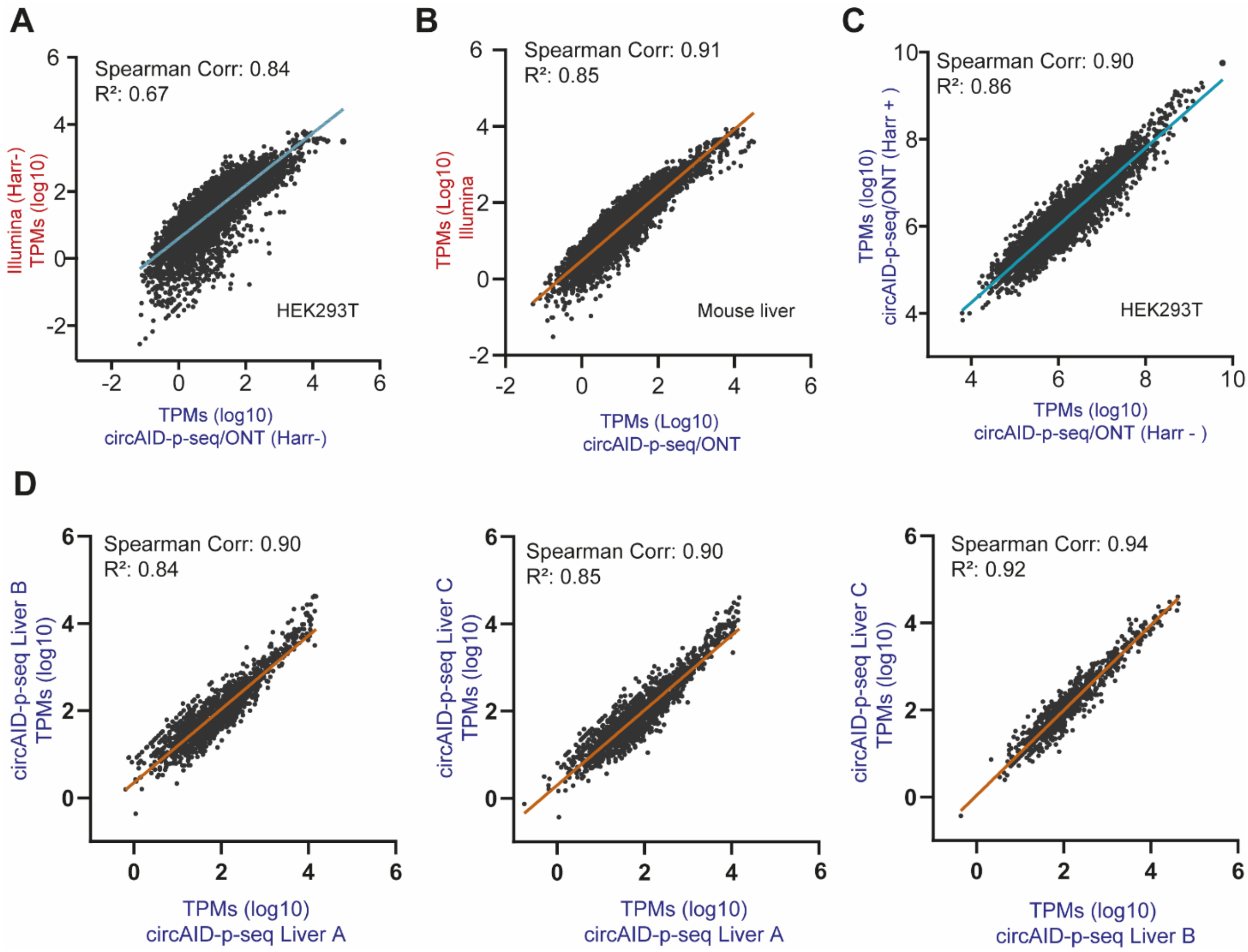


Figure S5. **Data correlation of Illumina and circAID-p-seq/ONT. (A)** RPF coverage correlation between circAID-p-seq/ONT and ILMN, in HEK293T experiment. Data represent only one replicate. Green line = linear regression **(B)** RPF coverage correlation between circAID-p-seq/ONT and ILMN, in mouse liver. Orange line = linear regression; data are mean of n= 3 biologically independent samples. **(C)** RPF coverage correlation between HEK293T treated and not treated with Harringtoine using circAID-p-seq/ONT method. Green line=linear regression. **(D)** RPF coverage correlation between the three biological replicates of mouse liver tissue (Liver A, B, C) using circAID-p-seq/ONT method. Orange line = linear regression**.** For all data, Spearman's rank correlation and squared Pearson linear correlation ($R^2$) are reported in figure. All genes are filtered for more than 1 count.

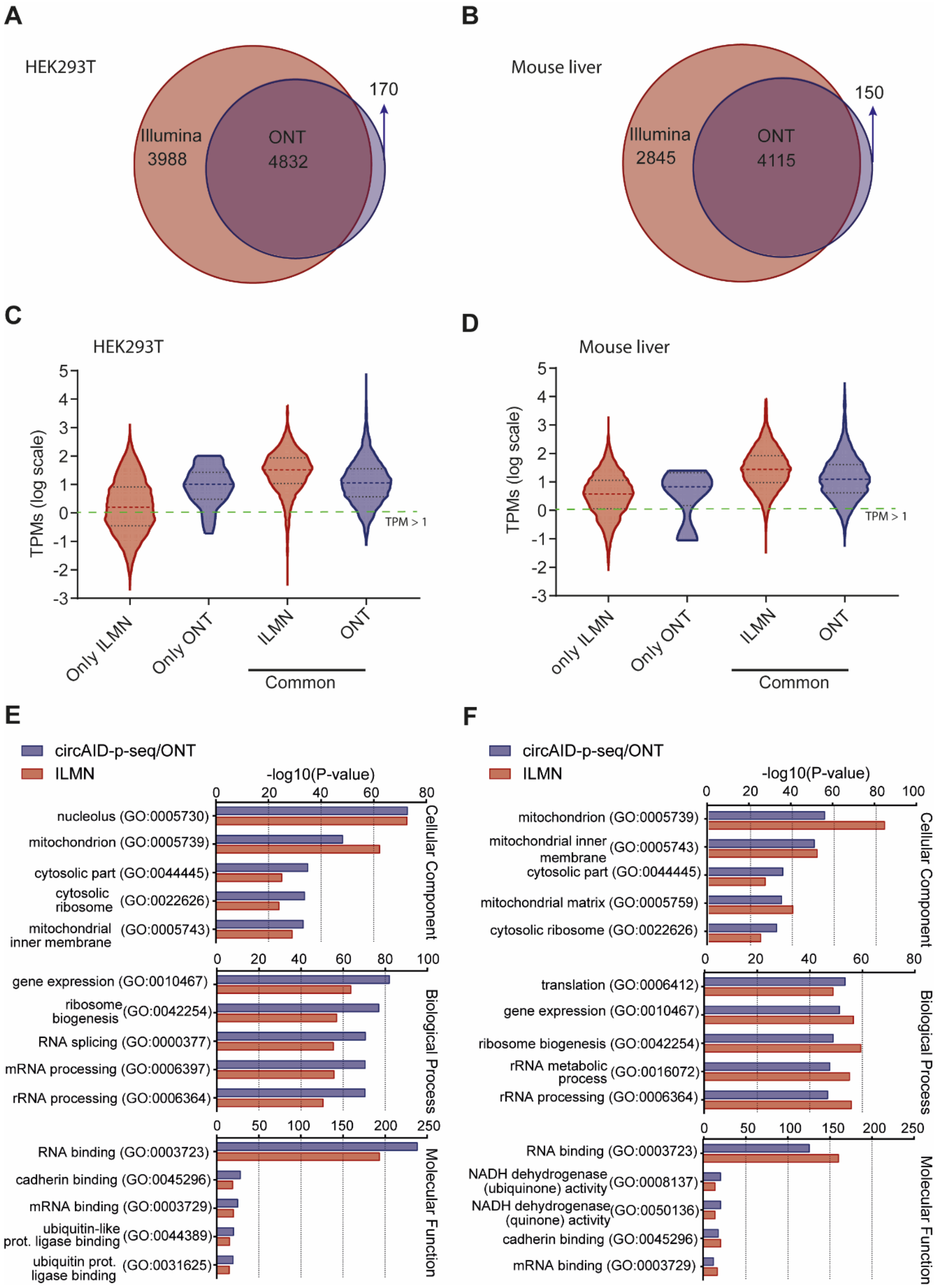


Figure S6. **Ribosome footprint data analysis. (A)-(B)** Venn diagram showing the number of common and unique genes (TPMs >10), between ILMN and circAID-p-seq/ONT, in (A) HEK293T and (B) mouse liver. **(C)-(D)** Violin plot showing the distribution of TPMs of genes identified (>1 count) with Illumina (ILMN) and circAID-p-seq/ONT (ONT), in (c) HEK293T and (**d**) liver mouse. Dashed line=median; dotted line=quartiles. For mouse liver tissues data are mean of n= 3 biologically independent samples. Green broken line, TPM > 1 threshold. **(E)-(F)** GO analysis for genes (TPMs >10) detected by circAID-p-seq/ONT (blue) and ILMN (red) in (E) HEK293 and (F) mouse liver. Top-5 enriched term for each category are sorted by value of –Log10(p-value).

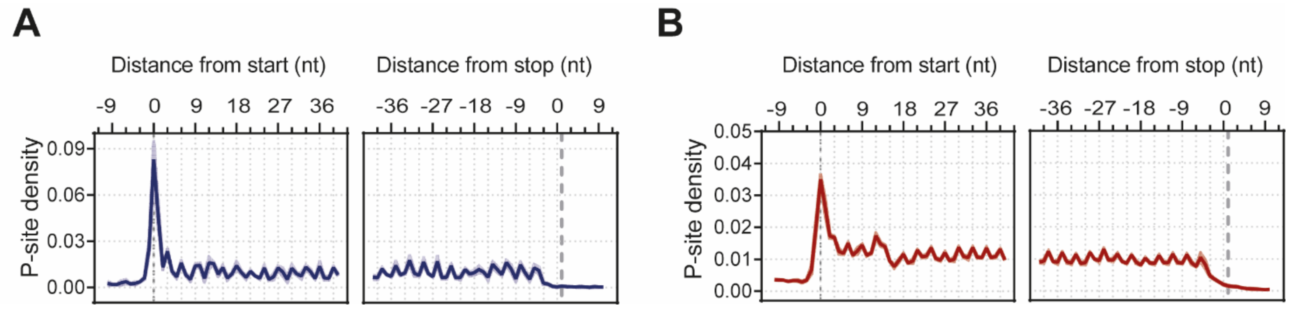


Figure S7. Metaprofiles for mouse liver tissue data showing the density of P-sites around translation initiation sites and translation termination sites for circAID-p-seq/ONT (**A**) and Riboseq/ILMN (**B**), using only common transcripts detected by both technology (n=4115; > 10 TPMs). Data are mean ± s.e.m. of n= 3 biologically independent samples (shadow).
